# Scalable Mesenchymal Stem Cells Enrichment from Bone Marrow Aspirate using Deterministic Lateral Displacement (DLD) Microfluidics Sorting

**DOI:** 10.1101/2023.05.03.539013

**Authors:** Nicholas Tan Kwan Zen, Kerwin Zeming Kwek, Teo Kim Leng, Mavis Loberas, Jialing Lee, Chin Ren Goh, Da Hou Yang, Steve Oh, James Hui Hoi Po, Simon M. Cool, Han Wei Hou, Jongyoon Han

**Author notes:** Co-First Authors (Nicholas Tan Kwan Zen, Kerwin Zeming Kwek).

## Abstract

The growing interest in regenerative medicine has opened new avenues for novel cell therapies using stem cells. Bone Marrow Aspirate (BMA) is an important source of stromal mesenchymal stem cells (MSCs). Conventional MSC harvesting from BMA relies on archaic centrifugation methods, often leading to poor yield due to osmotic stress, high centrifugation force, convoluted workflow, and long experimental time (∼ 2 – 3 hours). To address these issues, we have developed a scalable microfluidic technology based on Deterministic Lateral Displacement (DLD) for MSC isolation. This passive, label-free cell sorting method capitalizes on the morphological differences between MSCs and blood cells (leukocytes and RBCs) for effective separation using an inverted L-shaped pillar array. To improve throughput, we developed a novel portable multiplexed DLD system that can process 2.5 mL of raw BMA in 20 ± 5 minutes, achieving a 2-fold increase in MSC recovery compared to centrifugation methods. Taken together, we envision the developed DLD platform will enable fast and efficient isolation of MSCs from BMA for effective downstream cell therapy in clinical settings.

## I. Introduction

The advent of regenerative medicine marks a new paradigm shift in medical treatment through therapeutic cell delivery by autologous or allogenic means to replace/regenerate damaged cells, thereby creating personalized medicine^1, 2^. The discovery of stromal Mesenchymal Stem Cells (MSCs) has led to the pursuit of pioneering novel therapies due to its tri-lineage differentiation potential into mesodermal cell types *in vitro* (adipocytes, chondrocytes and osteocytes)^3^. This unique multilineage differentiation has spearheaded MSC therapies to treat health conditions such as neurological disorders^4, 5^, cardiac ischemia^6^, diabetes^7, 8^, and cartilage diseases^9^. Apart from their unique differentiation capabilities, MSCs are also known to exhibit immunomodulatory properties^10^, enabling expanded cell therapeutic applications. These cells can be found in various biological samples such as adipose tissues^11^, umbilical cord blood^12^ and salivary glands^13^. A traditional MSC harvesting site is aspirating from human bone marrow (BMA)^3, 14-16^ as it has a relatively high MSC quantity compared to other tissues. Despite the abundance of MSCs in the bone marrow, they only constitute 0.01% and 0.005% within the Bone Marrow Mono-Nucleated Cells (BMMNC) population for males and females, respectively^17^. This advocates a critical need for more effective and efficient MSC extraction from biological samples.

Advancements in cell sorting methods exploit cells’ biophysical and biochemical properties. Current cell sorting techniques such as Fluorescence-Activated Cell Sorting (FACS) and Magnetic-Activated Cell Sorting (MACS) leverage on antibody labeling targeted to cell surface markers^18, 19^. These techniques can reproduce extremely high sorting resolution but are limited by the high cost of antibodies, bulky machinery, and laborious sample preparation. They also require technical expertise and high operating pressure, which may influence MSC consistency and cell integrity^20^. Besides, it has been challenging to determine MSC-specific cell surface markers^21^. Antibody binding on cells may also affect their native biological or signaling pathways ^22^, thus unsuitable for downstream therapeutic purposes.

The conventional MSC isolation technique from BMA adopts a label-free density-based centrifugation^23-25^ method known as Ficoll Density Gradient Centrifugation (DGC). Through Ficoll DGC, the Mono-Nuclear Cell (MNC) fraction contains a heterogenous cell population consisting mainly of monocytes, lymphocytes, Hematopoietic Stem Cells (HSCs) and MSCs. Despite its wide usage, Ficoll DGC has the poorest yield amongst different MSC isolation techniques, including RBC lysis, immunomagnetic depletion, and simple filtration^26-28^. It was shown that Ficoll DGC typically suffers from significant cell loss amongst human progenitor cells (30% for HSC and 55% for MSC)^26^. MSCs can aggregate to form large clusters, congregating in the Polymorphonuclear (PMN) fraction during centrifugation^29^. Apart from the modest cell recovery, the workflow for Ficoll DGC (∼2 to 3 h) suffers from poor repeatability due to inconsistent preparation protocols, clinical personnel training, and increased exposure to contamination.

Microfluidic technologies have generated substantial advancement in cell sorting technologies over the years^20, 30^. Cell sorting using microfluidics often leverages the intrinsic biophysical properties of cells, rendering the techniques label-free^31^. They can be classified into two groups: active and passive methods. Active methods exploit externally applied fields or forces such as acoustophoresis^32^, electrokinetics^33^ and magnetophoresis^34^ to enhance separation. In contrast, passive methods rely on inherent hydrodynamic interactions between the channel structures and cells for separation, optimized by manipulating channel geometries^20^. Passive methods include inertial focusing^35^, Deterministic Lateral Displacement (DLD)^36^ and hydrodynamic filtration^37^. These microfluidic techniques are often easy to operate, integrate and scalable into clinical sample processing workflows. To our knowledge, no work has been done on directly isolating MSCs from whole undiluted BMA using microfluidic devices.

This study aims to determine the potential of employing DLD technology for direct BMA processing as an alternative to Ficoll DGC. We demonstrate a simple user–friendly workflow for BMA processing using a portable benchtop microfluidic pressure system. DLD, first described by Huang *et al*. ^36^ in 2004, leverages micropillar arrays to sort cells based on size and deformability with promising recovery rate and purity^38^. The sorting criteria (critical diameter, *D*_*C*_) can be calculated by an empirical formula based on the geometrical parameters of the pillar arrays where particles larger than the *D*_*C*_ are displaced laterally, away from their original stream^39^. Particles smaller than the *D*_*C*_ will remain in its original stream, causing a distinct separation between the larger and smaller cells. DLD sorting using peripheral blood samples has produced high recovery and purity of nucleated cells^40-42^. However, the cellular composition between peripheral blood and BMA differs. The latter comprises a higher amount of immature blood cells and nucleated cells with larger variations in cell biophysical properties, such as cell size, as studied by Xavier *et al*. ^43^. Thus, the efficiency of direct MSC isolation from undiluted BMA samples remains unknown.

Three parameters, namely recovery rate, total experimental time, and MSC Colony Forming Unit-Fibroblast (CFU-F) count validated through FACS analysis, will be used to determine the effectiveness and efficiency of MSC isolation between DLD sorting and Ficoll DGC. To improve throughput scalability, we also have developed a novel multiplexed DLD system that can process 2.5 mL of raw BMA in 20 ± 5 minutes, achieving a 2-fold increase in MSC recovery compared to Ficoll DGC protocol. Overall, the DLD platform shows excellent potential for fast and efficient isolation of MSCs from BMA in clinical settings.

## II. Methods and materials

### A. DLD device design

A multiplexed chip consisting of four DLD sorters (Figure 1b), connected in parallel, was developed to increase the processing speed for BMA. This multiplexed chip involves layering two PDMS substrates where the second PDMS layer comprises channel connectors combining four channels into one, thereby reducing experimental complexity. Each DLD device design is based on a mirrored DLD pillar array and a centric sheath flow. Samples are injected from the sides of the device, and particles larger than the *D*_*C*_ are displaced laterally into the center sheath stream and sorted into the collection outlet (Figure 2a). Particles smaller than the *D*_*C*_ follow its original stream, and they are not displaced, causing a separation between the smaller and larger particles. Quantification of cells in BMA from flow cytometry in previous studies^44-46^ provides little information on biophysical properties such as size, shape and deformability of MSCs in BMA. Thus, the effective *D*_*C*_ of the device used in this study is approximately 5.6 *μm* with an unconventional inverted L-shaped pillar array instead of the original circle pillar array. This DLD sorting specification was designed to negatively sort out the smaller and deformable red blood cells in BMA, which have been proven to work best using an inverted L-shaped pillar^38, 42^.

**Figure 1.**
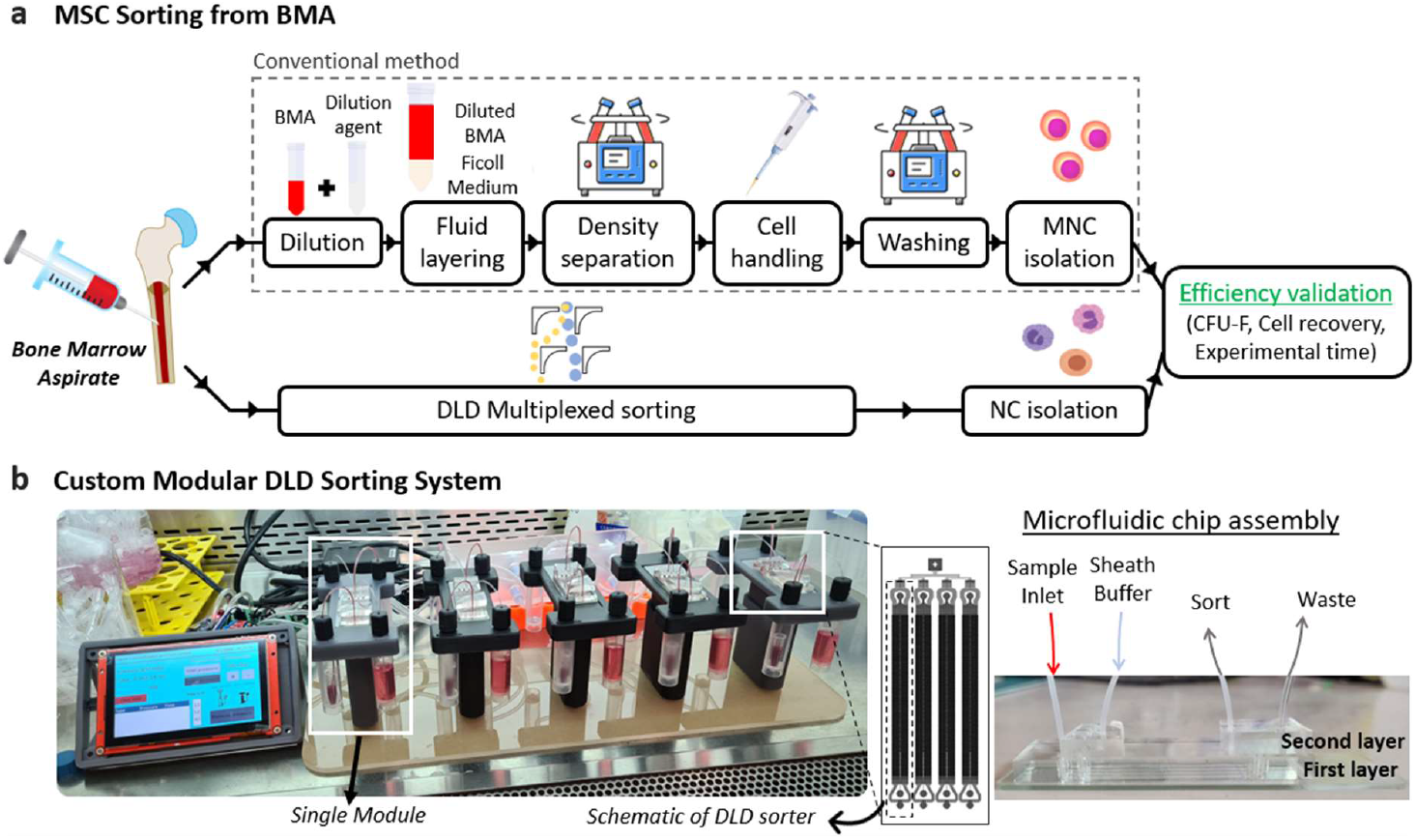
Schematics of the experimental comparison of DLD versus Ficoll DGC for MSC sorting and recovery from BM samples. (a) Experimental overview for bone marrow aspirate using Ficoll DGC and DLD sorting. The product from both methods will be subjected to MSC phenotypic assays to verify the presence of MSC through its colony forming capabilities (CFU-F). Cell recovery and experimental time will be used to corroborate the effectiveness and efficiency for both methods. (b) The multiplexed design parallelizes four DLD sorting array into a single chip to increase the sorting speed. Throughput is further increased using the custom sorting system which allows multiple chips (up to 5) to operate simultaneously.

**Figure 2.**
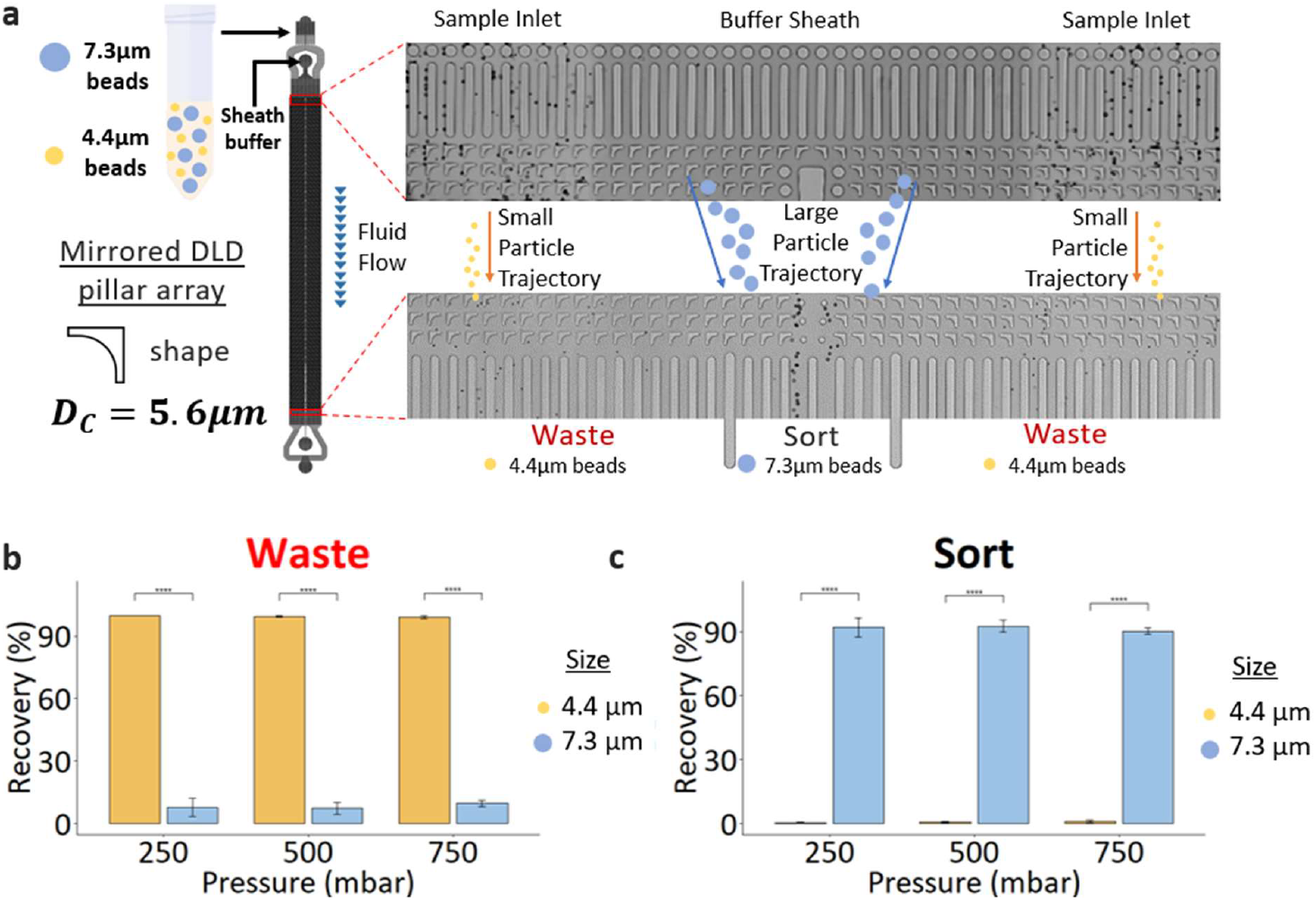
Characterisation of DLD sorting device. (a) Sorting characterisation of the DLD device using 4.4 μm and 7.3 μm beads. The mirrored DLD device consist of an input region which has a central sheath buffer flow sandwiched by two sample co-flow streams. The outlet region of the DLD device shows the sorted region with two side waste channels which is characterised by the 4.4 μm and 7.3 μm shown in the waste and outlet (sort) region respectively. (b) Graph showing the beads recovery in both the waste channels. (c) Graph showing the beads recovery in the sort channel. Paired two-tail student t-test was used to determine the statistical differences between outputs. **** denote p-value < 0.0001 for n = 4 samples.

### B. DLD device fabrication and preparation

Fabrication of a silicon master mold follows a standard photolithographic process using a chromed quartz photomask (JD Photodata, UK). A SU-8 mold was made using SU-8 2020 spun at 4500 RPM, resulting in a final resist thickness of 18 *μm*. The fabrication of the master mold was made using standard photolithography techniques via exposure, post-exposure bake and development. The master mold underwent a salinization step using Trichloro(1H,1H,2H,2H-perfluorooctyl)silane (Sigma-Aldrich, Singapore) to ensure a hydrophobic coating and ease of downstream DLD chip manufacturing. The production of DLD chips follows a soft lithographic process using the master mold. PDMS (Dow Corning, USA) is mixed in a 10:1 ratio with its curing agent and poured over the master mold taped onto a petri dish. The mixture will be degassed in a desiccator for 30 minutes before baking the degassed PDMS at 80°C for 2 hours. After curing, a surgical knife cuts the PDMS devices from the mold.

Holes of different sizes are punched at the ingress and egress of the desired chip (1*mm* for sample inlet, sheath flow and outlet (Sort), 1.5*mm* for waste). The resulting PDMS product will be bonded onto a glass slide cleaned with 70% ethanol, using oxygen plasma treatment and placed onto a hot plate at 140°C for 10 minutes to ensure complete bonding. PDMS-to-PDMS bonding follows the same protocol as PDMS-to-glass bonding, with the only difference being placing the bonded product into an oven at 80°C overnight instead of a hot plate.

Prior to experiments using biological samples, DLD chips are primed using 2% w/v Pluronic™ F-127 (Sigma, SG) with 20% Isopropyl alcohol (IPA) for 30 minutes to prevent nonspecific adsorption and easy bubbles removal due to its low surface tension properties from the ingress and 10 minutes from the egress using a pressure pump. After which, the chips will be washed with Dulbecco’s Phosphate-Buffered Saline (DPBS) (Thermo fisher scientific PTE LTD, Singapore) or Gibco Dulbecco’s Modified Eagle Medium (GE Healthcare, USA) for 5 minutes from the ingress to cultivate a suitable cellular environment as Pluronic™ F-127 is toxic to cells.

### C. Single-device characterization

To characterize sorting performance, the DLD chip is placed onto a microfluidic device platform with imaging capabilities powered by an Arduino microcontroller and a positive displacement pump (RS PRO, Singapore). Custom fittings are designed and 3D-printed using low-force stereolithography (Form 3, Formlabs), forming an airtight junction between the sample reservoir and tubing (Cole-Parmer, USA) using O-rings. The tubing length stands at 30 *mm*. Microsphere counting techniques leverage computer vision technology, and cell counting is performed using a hemocytometer.

### D. Modular high-throughput sorting

For high throughput sorting targeted for clinical applications, a new concise portable benchtop multiplexed microfluidic platform was developed, which allows multiple chips to operate concurrently. The design aims to achieve portability, robustness, and simplicity so that non-trained users can operate the sorter. We have also established a clear, easy-to-follow experimental workflow to complement our system. Tubings, fittings, and the prototype are sterilized in a Biological Safety Cabinet (BSC) under UV light for 30 minutes before use. Experiments using BMA are also performed in the BSC to ensure sterility.

A pressure sensor (AMS 5915, AMSYS, Germany) and pump setup were assembled to control the desired operating pressure. To offset the pressure drop at the tubings and connections, we integrated a simple pressure cutoff control in our PCB design that monitors the pressure continuously and activates the pump whenever the sensor detects a drop below the 10% threshold. Pneumatic solenoid valves (Parker Hannifin, USA) were connected to a manifold inside the pressure chamber to enable proper distribution of pressure to the correct ports. The pressure chamber with manifold, modular unit, and Human-Machine Interface (HMI) stand is fabricated using 3D printing.

### E. Peripheral blood samples

Peripheral whole blood from healthy donors validated the DLD device for extracting nucleated cells from RBCs and platelets. 3 mL of blood per donor was stored in a 3 mL EDTA tube. The blood sample was processed within 4 hours of the blood draw to prevent inherent aggregation and degradation of samples. The blood samples were stored at room temperature to ensure minimal biophysical changes to the nucleated cells.

### F. Bone marrow aspirate samples

Approximately 50 mL of bone marrow was harvested from the iliac crest of three healthy donors aged 26 – 52 under local anesthesia, following informed consent and strict adherence to ethical guidelines (IRB 2019/00284). The aspiration was performed using a Jamshidi needle, collected into heparinized syringes, and transferred to sterile containers. The bone marrow aspirate was transported to the National University of Singapore Tissue Engineering Programme (NUSTEP) for further processing. From the 50mL aspirated, 5 mL was used for Ficoll DGC and DLD sorting at the Bioprocessing Technology Institute, A*STAR, Singapore. The remaining aspirate was not used in this study.

### G. BMA processing using Ficoll DGC

Heparinized bone marrow aspirate was first diluted with an equal amount of D10 culture media. This culture media consisted of Dulbecco’s Modified Eagle’s Medium (DMEM), 1,000 mg/ml glucose supplemented with 10% fetal bovine serum and 2 mM L-glutamine. The diluted aspirate is carefully layered on top of 1.078 g/ml Ficoll-Paque PREMIUM (Cytiva, USA) at a ratio of 2.5 mL diluted aspirate to 2 mL Ficoll-Paque. The solution was centrifuged at 400 g for 30 minutes at ambient temperature without brake mode. Next, the interface layer containing the mononuclear cells was collected and washed twice by centrifugation at 200 g for 15 minutes at maximum brake mode using complete DMEM.

### H. Mononuclear cell culture

Following the user guide, cells obtained from the Ficoll DGC method and DLD sorting were counted using Nucleocounter NC-250 (Chemometec, Denmark). Cells (3.5 × 10^6^) were seeded into a 25 cm^3^ tissue culture flask (ThermoFisher Scientific, Singapore) with 5 mL of D10 culture media. DNase I (Roche, Germany) was added into the culture media at 20 units/mL concentration. Culture media change was done every three to four days intervals. The spent culture media was centrifuged at 380 RCF for 5 minutes at each culture media change. The supernatant was discarded, and the pellet was resuspended in 5 mL fresh culture media. DNase I at 20 units/mL was added. The fresh media with resuspended cells was seeded into the cell culture. Cells were cultured until 80% confluency was reached. These cells are termed Passage 0 mesenchymal stem cells (P0 MSC).

### I. Mesenchymal stem cell culture

Confluent P0 MSC cultured in 25 cm^3^ tissue culture flasks were passaged using 0.25% Trypsin-EDTA dissociation solution (Gibco, Singapore). The passaged cells were counted using Nucleocounter NC-250 and seeded into a 75 cm^3^ tissue culture flask (ThermoFisher Scientific, Singapore) with 15 ml of D10 culture media at a seeding density of 5000 cells/cm^3^. Cells were cultured until 80% confluency was reached. These cells are called Passage 1 mesenchymal stem cells (P1 MSC) at this stage. These P1 MSCs were then further passaged and seeded into (i) 175 cm^3^ tissue culture flask (ThermoFisher Scientific, Singapore) with 35 mL D10 culture media at a seeding density of 5000 cells/cm^3^ and cultured until 80% confluent, and (ii) 6-well tissue culture plate (ThermoFisher Scientific, Singapore) with 2 mL D10 culture media at a seeding density of 65 cells per well and cultured for 14 days with culture media change on Day 7 for CFU-F analysis. These cells were termed Passage 2 mesenchymal stem cells (P2 MSC).

### J. MSC quality analysis: Colony Forming Unit – Fibroblast (CFU-F)

P2 MSCs cultured on a 6-well tissue culture plate at a seeding density of 65 cells per well for 14 days with culture media change on Day 7 were used for CFU-F analysis^47^. The cells were washed with sterile Phosphate Buffered Saline (PBS) (Gibco, Singapore) and stained with 2 mL of CFU-F stain for 30 minutes at room temperature. The CFU-F stain was prepared by dissolving a 0.5% weight-to-volume ratio of Crystal Violet (Sigma-Aldrich, Singapore) in a 25% volume-to-volume ratio of Methanol (Sigma-Aldrich, Singapore) in deionized water. The cells were then rinsed with PBS thrice to remove excess stain. The plate containing the stained cells was dried. Once the plate had dried, the number of colonies formed was counted. A colony is defined as a region with a minimum of 50 cells. Any areas with less than 50 cells were not considered a colony and, thus, were not counted.

### K. MSC quality analysis: Cell Surface Marker (FACS)

P2 MSCs cultured in a 175 cm^3^ tissue culture flask at a seeding density of 5000 cells/cm^3^ until 80% confluency were used for cell surface marker (FACS) analysis. Firstly, cells were washed with sterile PBS, dissociated from the tissue culture flask using 0.25% Trypsin-EDTA, and counted using Nucleocounter NC250. 1 × 10^5^ cells were used for each cell surface marker analysis. The positive selection markers used for the study were CD73, CD90, CD105, and CD146, and the negative selection markers were CD34 and CD45. Cells were washed in FACS buffer, which was made by dissolving a 1% weight-to-volume ratio of Bovine Serum Albumin (Sigma-Aldrich, Singapore) in PBS. The washed cells were then added to a 96-well V-bottom plate (Greiner Bio-One, Germany) and individually incubated with each fluorophore-conjugated antibody of positive and negative selection markers (Biolegend, USA) on ice for 30 minutes. The cells were washed with FACS buffer to remove excess unbound antibodies and resuspended in FACS buffer. According to the user manual, cells were analyzed using Novocyte (ACEA Biosciences, USA).

## III. Results

### A. DLD sorting characterization and optimization

To calibrate the DLD chip, polystyrene microspheres (Bangs Laboratory, USA) of sizes 4.4 *μm* and 7.3 *μm* were used at a stock 1.125% w/v concentration. This test illustrates the size-based sorting capabilities as the rigid beads are larger and smaller than the *D*_*C*_ (5.6 *μm*). As the pressure and corresponding flow rate increased, the sorting of 7.3 *μm* beads showed a minor decrease in beads recovery (92.14 ± 3.85% to 90.25 ± 1.52%). Using the inverse L-shaped pillars could result in possible anisotropic effects shown by Vernekar *et al*^*48*^. By comparing the sorting efficiency of 4.4 *μm* beads, the DLD device with a *D*_*C*_ of 5.6 *μm* performs as designed in the operating pressure range of 250 mBar to 750 mBar. The 7.3 *μm* beads accounted for 7-10 % of the total count at the waste outlet (Figure 2b) and were not deflected via DLD (See Supplementary Video S1 and S2). This is likely due to the functional design of the device, where the inlet sample flow interacts with the DLD array side walls resulting in inefficient sort near the walls.

### B. Peripheral blood sorting

Peripheral blood was used to elucidate the cell deformability effects and sorting efficiency of the DLD device. Unlike rigid beads, it is known that cell deformability impacts the DLD sorting efficiencies^41, 43, 49, 50^. The same DLD device used in Figure 2 was scaled linearly up to 4x by connecting the devices in parallel with a second layer microfluidic sample distribution channel at the input and output. Undiluted and unfiltered whole blood was processed using the device, as shown in Figure 3a and Supplementary Video S3.

**Figure 3.**
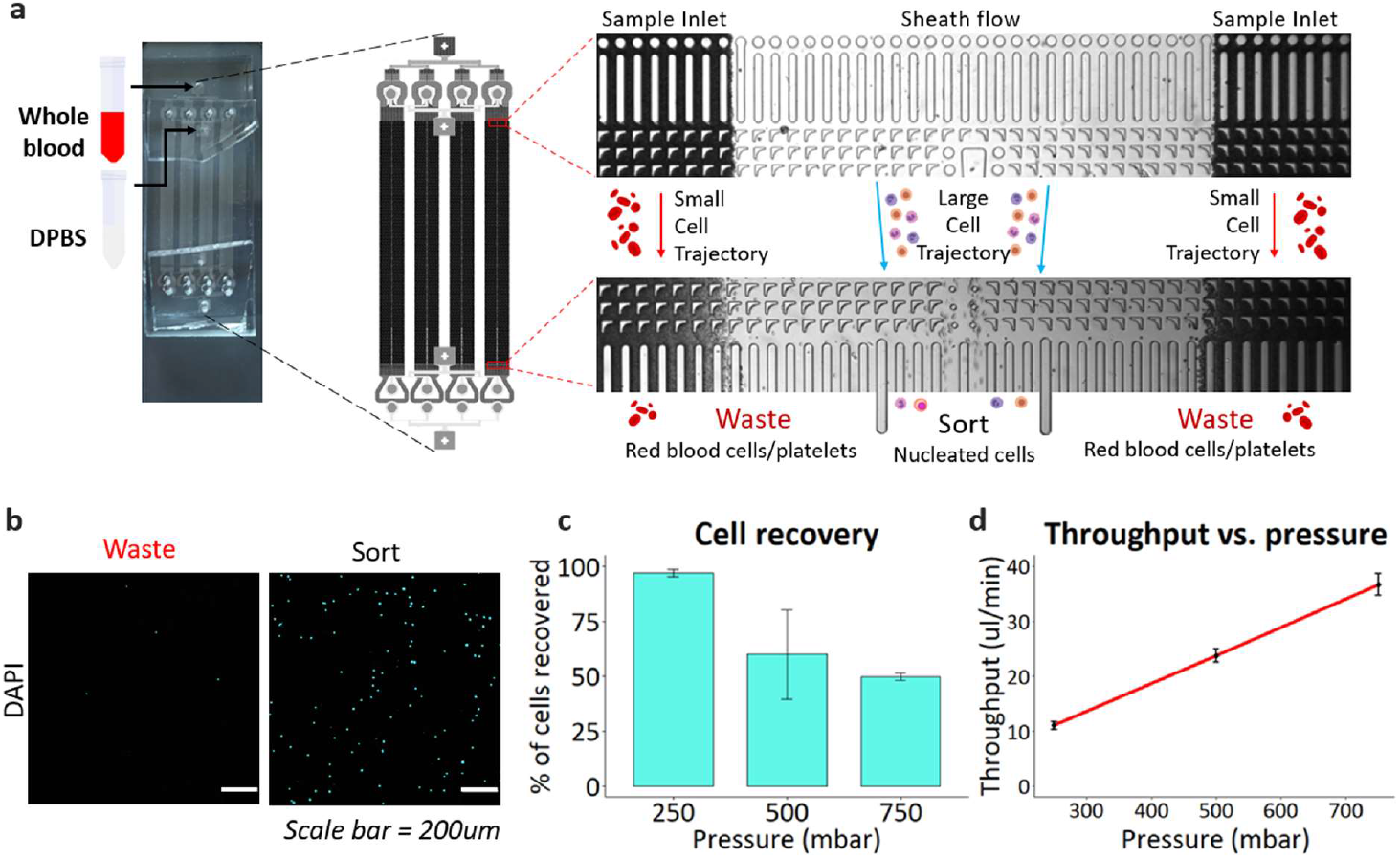
Characterizing whole blood nucleated cell sorting in parallel DLD devices. (a) PDMS DLD device shown with 4x parallel devices to increase sorting throughput. The input and output region of the DLD device can be seen with the sample stream co-flow with a sheath buffer stream. Nucleated cells were collected in the ‘sort’ stream while the waste stream consists RBCs and platelets. (b) Fluorescence images showing the nucleated cells stained with Hoechst 33342 for counting of nucleated cells in the respective sorted outlets at 250 mbar. (c) Cell recovery from the DLD device was measured with respect to varying flowrates. (d) Graph showcasing the corresponding flowrates (ul/min) and pressures used (mbar) for a fixed volume (100 uL).

To facilitate the output cell count, cells are DNA stained with Hoechst 33342 fluorescent stain solution and counted with a hemocytometer using Brightfield and DAPI microscopy. With fluorescent field imaging, the cell recovery rate is quantified through manual counting and carried out as the following:

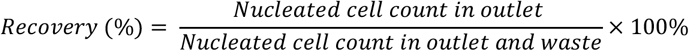

The corresponding results (Figure 3d) showed cell recovery decreases as pressure increases. Theoretically, the DLD device design with *D*_*C*_ = 5.6 *μm* should result in a 100% efficiency for both sorted (PBMC size > 7.23 ± 0.3 *μm*)^51^ and unsorted products (RBC size∼3 *μm*, platelets size = 3.64 ± 0.72 *μm*)^52, 53^. However, in comparison to the beads sorting, peripheral blood results showed different efficiencies across different pressure values, which underscore the need to account for an additional cell deformability factor. Decoupling size and deformability effects in DLD sorting remain challenging, and thus, various groups often use the effective size parameter to quantify the deformed size of a biological cell^49, 50, 54^. This observation for stem cell sorting is made following studies done by Xavier et al, where they verified the deformability effects in DLD sorting using MG-63 and HL-60 cells^43^ with varying flow rates. Therefore, at higher flow rates, the increased forces acting upon the cell induce a larger change in size as compared to the literature, which explains the decrease in efficiency (Figure 3d). However, we have decided to use the highest pressure for our BMA experiments for two reasons. The trade-off between high throughput and cell yield must be considered based on different applications. For this study, achieving high throughput is critical for manufacturing-scale cell separation and favours higher flow and volume processing rates. Secondly, peripheral blood is also known to express pseudoplastic properties^55^. Since viscosity is inversely proportional to flow rate, we would want blood/BMA to exhibit less viscosity at a higher flow rate, which is beneficial to achieve maximum volume processing rates. In this case, we assume that the properties of BMA and peripheral blood are similar due to the abundance of RBCs.

### C. MSC isolation from Human Bone Marrow Aspirate using DLD

To validate our proof of concept that DLD sorting works better than Ficoll DGC, we performed two comparative sorting experiments: Ficoll centrifugation and DLD sorting. The Ficoll DGC procedure has been explained in section 2G. To preserve physiological relevance, undiluted human BMA was processed for DLD sorting. As such, the resulting cell recovery and the number of clonogenic cells are expected to be lower than in other studies, where cell separation is optimized through additional steps such as dilution, RBC lysis, and selective cell combination^26, 27, 43, 56, 57^. However, because the primary goal of this study is to introduce a simple, single-step cell extraction protocol, such an experimental design can be justified. As part of efforts to increase throughput, five high-throughput chips were used, resulting in twenty parallel DLD devices operating concurrently.

The workflow is described in Figure 4. Firstly, we connected the tubings from the reservoir vials (sample fluid, sheath buffers, outlet (sort) and waste) to the chip on the modular platform. We then prime the device with the priming agent and wash chips with DPBS before flushing the device with DMEM. To start sorting, we loaded the undiluted BMA into the sample inlet port and entered the desired pressure into the HMI. Our undiluted BMA sample had a cell initial concentration of 8.59 ± 3 × 10^6^ *cells*/*mL*, and was subjected to a 20 *μm* filter before sorting to remove unwanted coagulations (see Supplementary Figure 1). After sorting, we combined the outlet (sort) vials from each module into a 15 mL vial before data analysis. This was done likewise for the waste vials. The measured cell recovery between peripheral blood and BMA at 750 mbar differ slightly (49.8 ± 1% and 43.9 ± 6.3%, respectively). Since the design of our DLD array is based on the static cell size, this illustrates the difference between nucleated cell sizes from BMA and peripheral blood, which is also noted in literature^45, 46^. One of the reasons for the size disparity could be due to interactions with the DLD pillars causing cells to deform and change in size. Despite the lower cell recovery rate, the yield is better than Ficoll DGC by a factor of two (Figure 5a). The time taken for DLD to process 2.5 mL is within minutes and can be improved by increasing the number of modules (more chips operating concurrently).

**Figure 4.**
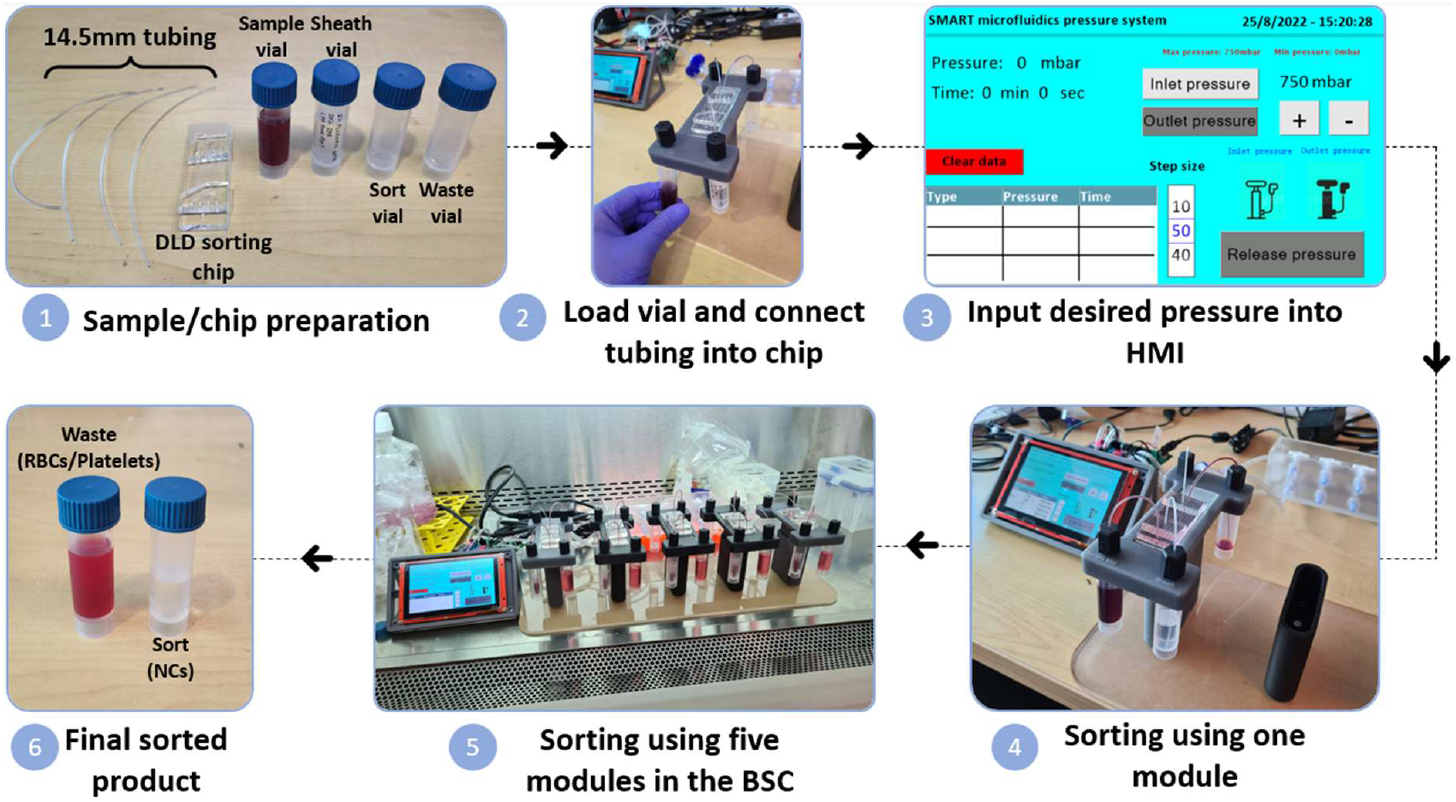
Experimental procedure and engineered system for scaled sorting operations of the DLD sorting microfluidic device(s). Step procedures for high throughput sorting using our in-house microfluidics pressure system with modular platforms. This 5 steps approach will allow for reduced user training and result variability associated with human error present in the Ficoll DGC protocol where the process is highly dependent on the user’s technical knowledge and experience.

**Figure 5.**
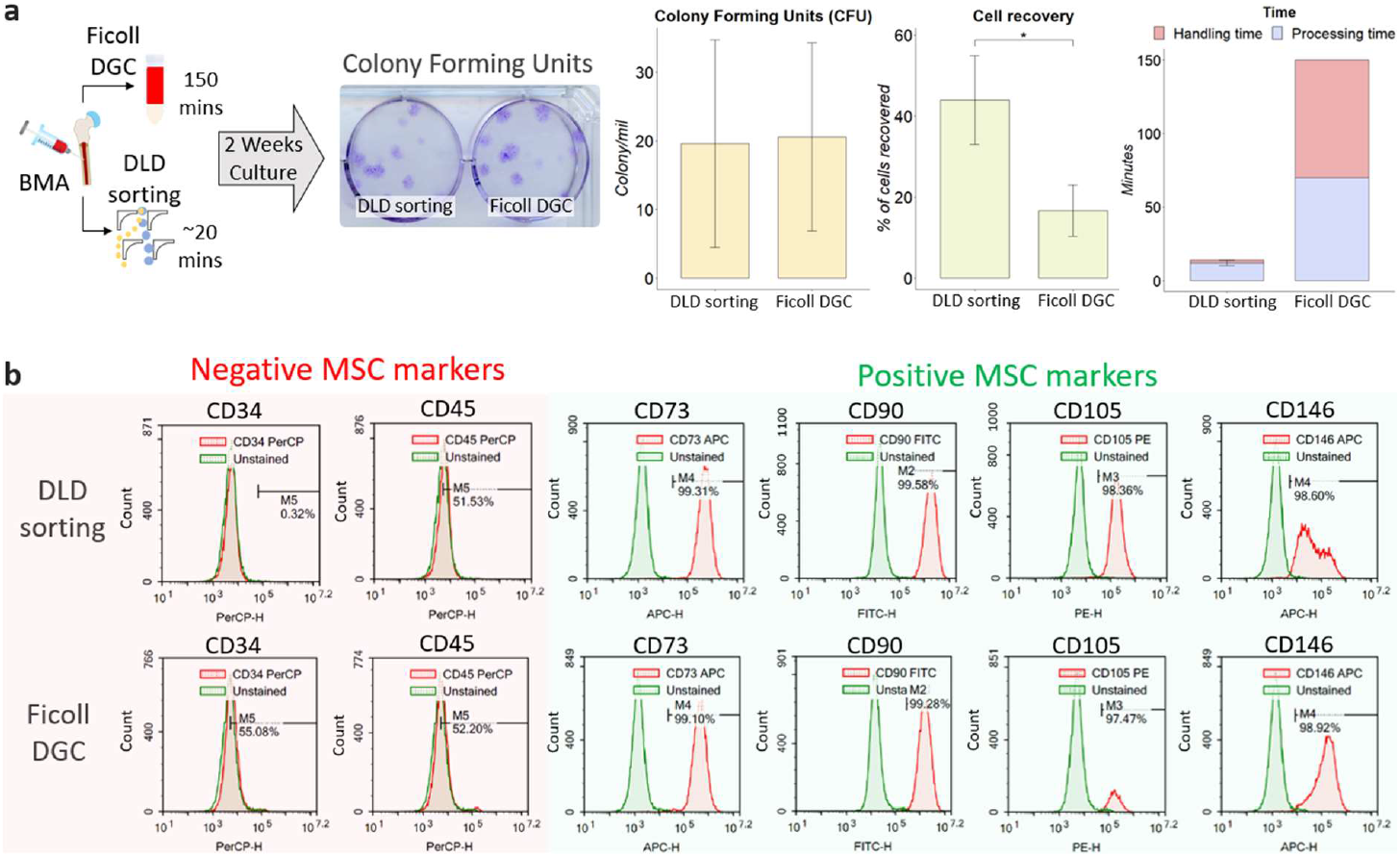
Sorting efficiency and recovery of MSC from BM samples using DLD device and Ficoll DGC. (a) BMA processing using both DLD sorting and Ficoll DGC was performed to validate the effectiveness of MSC enrichment through CFU count, cell recovery and time taken (n = 3). (b) Flow cytometry analysis of MSC phenotypic cell surface markers on sorted cells using DLD and Ficoll DCG.

In contrast, Ficoll DGC has a fixed, established protocol, and the time taken is independent of sample volume. The Ficoll DGC method can take 2 – 3 hours (volume processed can range from 2mL – 35mL with dilution), and a series of centrifugation steps are needed to achieve better results. The time plot in Figure 5a includes both processing and handling time. Processing time refers to the total sorting/isolation duration, and handling time refers to the time taken to handle the cells before/during the sorting, which includes washing and dilution. Studies have also shown that the Ficoll medium is toxic to cells^58^, and multiple centrifugations might increase cell death due to prolonged exposure to the Ficoll medium. The results showed that DLD sorting reproduced a comparable number of colonies per million cells to Ficoll DGC (Figure 5a). However, since the cell recovery is two times higher than Ficoll DGC, the total number of colonies and MSC recovery is significantly higher for DLD sorting. To verify that these colonies are MSCs, we performed a phenotypic assay after the second subcultivation (Passage 2) to examine for MSC surface marker expression (CD73, CD90, CD105, CD146) and clear of the hematopoietic cell lineage (CD34^−^, CD45^−^) using FACS as iterated under the section 2K. The findings (Figure 5b) indicate that the MSC population is similar in the Ficoll DGC colonies. However, the total number of MSCs is still higher in DLD sorting due to higher cell yield.

## IV. Discussion

Stem cell therapy is the new disruptive innovation in regenerative medicine. Multiple biological tissues have been reported to contain MSCs, and identifying the most effective MSC isolation technique from a suitable source is critical for sorting functional MSC. Despite having a low MSC yield, Ficoll DGC has long been the primary method for MSC isolation from BMA (Table 1).

**Table 1.**
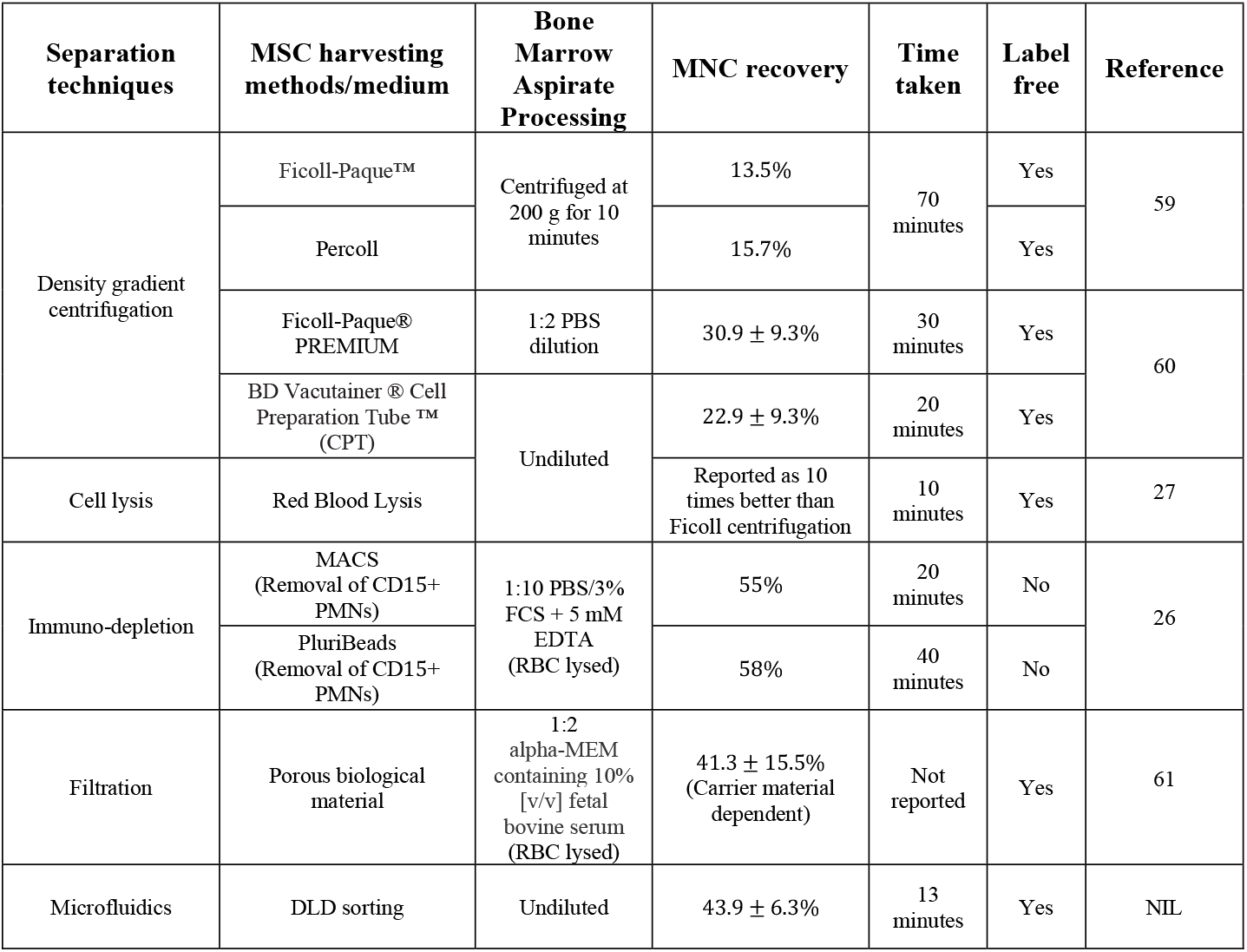
Comparison of MSC sorting techniques from BMA

Comparing the recovery across all MSC harvesting methods, our current DLD design offers one of the best alternatives with no sample dilution or manual washing steps (handling time). “Time taken” in Table 1. refers to the reported processing time of these methods and does not include sample handling time such as pipetting, centrifuge braking or sample filtration which could plausibly increase the experimental timing significantly. However, what sets the DLD sorting technology apart from the rest of these methods is the absence of sample preparation, pre and post-sort washing steps and potential scalability. The precision and continuous flow single-step sorting also eliminates variability associated with the cell handling steps. For the multistep Ficoll DGC process, the same personnel performing the technique will exhibit some form of variability due to human error, which is almost impossible to eliminate. The DLD sorting also inherently integrates a washing step where sorted cells are migrated into the buffer stream, thus removing the need for additional washing steps.

The current sorting efficiency for MSCs is 43.9 ± 6.3%, and improvements on the nucleated cell recovery and time can be achieved by further optimising microfluidic DLD specifications by reducing the critical size specification to sort out smaller ones or more deformable cells at the same or higher flowrate. It can be hypothesized that sorting smaller nucleated (mononuclear and polynuclear) cells will contain a higher yield of MSCs than only sorting mononuclear cells via Ficoll DGC. Early evidence from this paper has shown that mononuclear cell sorting loses a large fraction of MSCs. This was also validated by Horn *et al*^27^ employing whole blood lysis protocol to extract all nucleated cells showing that there exists an MSCs population that is missed out by mononuclear cell sorting methods. Given the similar cell recovery values between peripheral blood and BMA at 750mbar, we are confident that our DLD device can achieve a higher cell recovery (∼80% – 90%) if deformability effects are negated (running at 250mbar) at the expense of throughput. Also, it is often necessary to expand these cells to the desired cell quantity to meet the requirements for MSC-based therapies. However, increased passaging is known to impair MSC plasticity and contribute to genomic instability, telomere shortening and heterogeneity^62^, which limits the therapeutic effects of MSCs, thereby reinforcing the need for an effective sorting method. Whilst the CFU potential is comparable between the two approaches, DLD offers a higher cell recovery, translating to a higher MSC recovery. Given the rare occurrence of MSC in BMA, a higher yield during sorting can lead to exponential cell numbers in downstream expansion.

While microfluidic chip stacking has been explored by various groups^63, 64^ to enable scaling of sample processing, this paper investigates the possibility of scaling the sorting process using a modular sort unit that can be seamlessly integrated into a central pressure distribution setup. The microfluidic setup and processing workflow can also be used for leukapheresis with very high yield at lower flow rates. Without the need for multiple washing steps (Table 1.), the major benefit of using the DLD sorting technique is integrating bioreactors in closed-loop processing of cell separation and manufacturing with the potential to reduce contamination, workforce cost and improve consistency.

Isolating cells by label-free parameters such as size and deformability enables the enrichment of specific cell functions or phenotypes. The next questions to be answered following the sorting are: Do the sorted cells have differing growth potentials? What is the differentiation potential of MSCs to be of a particular phenotype for regenerative medicine? To quantify the qualities of these DLD-sorted MSCs, it is paramount that stemness and immunomodulation are validated through functional assays such as RT-PCR, which serves as a motivation for future study. A previous study by Yin Lu *et al* has ascertained that MSCs of sizes 17*μm* − 21*μm* have the most potential for chondrogenesis using inertial sorting^65^. While downstream quantitative assays to test for the mesoderm cell lineage differentiation were not explored in this study, the utility of DLD sorting goes beyond a sample processing tool with the potential for specific phenotypical enrichment for more efficacious cell manufacturing.

## V. Conclusion

In this study, we have developed a multiplexed DLD microfluidic chip for MSCs sorting from BMA samples directly and demonstrated superior separation performance compared to Ficoll DGC (conventional standard). Improving the device throughput and automation in our prototype makes this clinical workflow faster (∼20 min), more user-friendly and more robust for MSC isolation. With its label-free approach, this platform technology is low-cost, and we can envision it to be readily adapted to sorting other cell types by controlling DLD pillar shapes and geometries.

## Author Contributions

N.T.K.Z., K.K.Z., H.W.H., S.O., S.M.C., J.H. designed aspects of the project study. N.T.K.Z. and K.K.Z. designed and fabricated the chip. N.T.K.Z., K.K.Z, G.C.R., Y.D.H. optimized and performed the experiments. N.T.K.Z., K.K.Z. and G.C.R. designed, fabricated, and calibrated the electronics and sorting setup. N.T.K.Z. and K.K.Z. analysed the data. N.T.K.Z. coded the algorithms for software and hardware. J.H.H.P. performed the bone marrow aspiration. T.K.L., M.L., S.O. performed the patient recruitment and post-sort cell measurements and FACS profiling. N.T.K.Z., K.K.Z., S.M.C., H.W.H. and J.H. wrote the manuscript. All authors reviewed and approved of the manuscript prior to submission.

## Conflict of Interest

We declare no conflict of interest.

## Acknowledgements

We thank Afizah Binte Mohd Hassan and Ren Xiafei for ensuring seamless coordination and operations during patient recruitment. This research is supported by the National Research Foundation, Prime Minister’s Office, Singapore under its Campus for Research Excellence and Technological Enterprise (CREATE) programme, through Singapore-MIT Alliance for Research and Technology (SMART): Critical Analytics for Manufacturing Personalized-Medicine (CAMP) Inter-Disciplinary Research Group.

## Notes

### Competing Interest Statement

The authors have declared no competing interest.

