## Supplementary figures and images for "Scalable Mesenchymal Stem Cells Enrichment from Bone Marrow Aspirate using Deterministic Lateral Displacement (DLD) Microfluidics Sorting"

### Supplementary figure

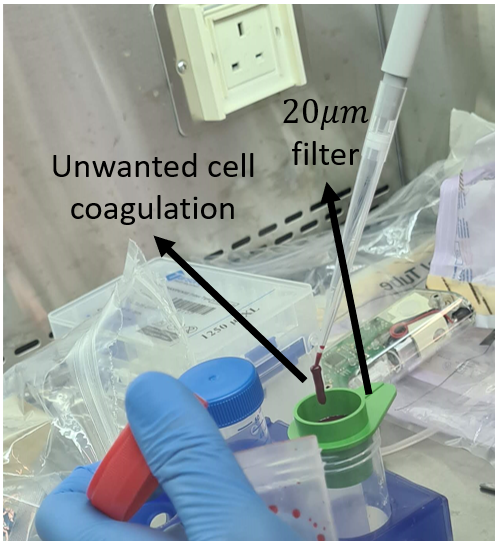

**Fig. S1** Cell coagulation can cause a clog in the device.
